# Gata4 drives Hh-signaling for second heart field migration and outflow tract development

**DOI:** 10.1101/427336

**Authors:** Jielin Liu, Henghui Cheng, Menglan Xiang, Lun Zhou, Ke Zhang, Ivan P. Moskowitz, Linglin Xie

## Abstract

Dominant mutations of Gata4, an essential cardiogenic transcription factor (TF), cause outflow tract (OFT) defects in both human and mouse. We investigated the molecular mechanism underlying this requirement. Gata4 happloinsufficiency in mice caused OFT defects including double outlet right ventricle (DORV) and conal ventricular septum defects (VSDs). We found that Gata4 is required within Hedgehog (Hh)-receiving second heart field (SHF) progenitors for normal OFT alignment. Increased Pten-mediated cell-cycle transition, rescued atrial septal defects but not OFT defects in Gata4 heterozygotes. SHF Hh-receiving cells failed to migrate properly into the proximal OFT cushion in Gata4 heterozygote embryos. We find that Hh signaling and Gata4 genetically interact for OFT development. Gata4 and Smo double heterozygotes displayed more severe OFT abnormalities including persistent truncus arteriosus (PTA) whereas restoration of Hedgehog signaling rescued OFT defects in Gata4-mutant mice. In addition, enhanced expression of the Gata6 was observed in the SHF of the Gata4 heterozygotes. These results suggested a SHF regulatory network comprising of Gata4, Gata6 and Hh-signaling for OFT development. This study indicates that *Gata4* potentiation of Hh signaling is a general feature of *Gata4*-mediated cardiac morphogenesis and provides a model for the molecular basis of CHD caused by dominant transcription factor mutations.

**Author Summary:** Gata4 is an important protein that controls the development of the heart. Human who possess a single copy of Gata4 mutation display congenital heart defects (CHD), including the double outlet right ventricle (DORV). DORV is an alignment problem in which both the Aorta and Pulmonary Artery originate from the right ventricle, instead of originating from the left and the right ventricles, respectively. To study how Gata4 mutation causes DORV, we used a Gata4 mutant mouse model, which displays DORV. We showed that Gata4 is required in the cardiac precursor cells for the normal alignment of the great arteries. Although Gata4 mutation inhibits the rapid increase in number of the cardiac precursor cells, rescuing this defects does not recover the normal alignment of the great arteries. In addition, there is a movement problem of the cardiac precursor cells when migrating toward the great arteries during development. We further showed that a specific molecular signaling, Hh-signaling, is responsible to the Gata4 action in the cardiac precursor cells. Importantly, over-activating the Hh-signaling rescues the DORV in the Gata4 mutant embryos. This study provides an explanation for the ontogeny of CHD.

## Introduction

Congenital Heart Defects (CHDs) CHDs occurr in approximately 1% of live births [1] and are the most common serious birth defects in humans [2, 3]. Approximately one third of the CHDs involve malformations of the outflow tract (OFT), which leads to significant morbidity and mortality of children and adults [4]. Multiple OFT abnormalities involve the relationship of the Aorta and Pulmonary Artery to the underlying left and right ventricles. For example, double-outlet right ventricle (DORV) is an anomaly in which the Aorta and Pulmonary Artery originate from the right ventricle [4]. A key characteristic of DORV that distinguishes it from other OFT defects is that the aorta and pulmonary trunk are well separated but are improperly aligned over the right ventricle. The molecular basis of OFT misalignment in DORV has remained unclear.

SHF-derived cells migrate into the developing poles of the heart tube, to effect morphogenesis of the cardia cinflow and outflow. The anterior SHF is essential for OFT and great artery development [5-9]. The failure of the anterior SHF-derived myocardial and endocardial contributions to the arterial pole of the heart causes a shortened OFT and arterial pole misalignment, resulting in inappropriate connections of the great arteries to the ventricular mass [10-12]. Deletion of genes responsible for SHF morphogenesis, such as *Isl1*, *Mef2c*, and *Jagged1*, leads to abnormal OFT formation including DORV [5, 6, 8, 12-19]. These observations lay the groundwork for investigating the molecular pathways required for OFT development in SHF cardiac progenitor cells.

Gata4, a member of the GATA family of zinc finger transcription factors, is an essential cardiogenic transcriptional regulator implicated in many aspects of cardiac development and function [20-34]. Human genetic studies have implicated haploinsufficiency of GATA4 in human CHDs, to date including atrial septal defects (ASD), ventral septal defects (VSD), and tetralogy of Fallot (TOF) [21, 35-39]. In mouse models, decreased expression of *Gata4* results in the development of common atrioventricular canal (CAVC), DORV, and hypoplastic ventricular myocardium in a large proportion of mouse embryos [27, 40]. Multiple studies have demonstrated the molecular requirement of Gata4 in the endocardium for normal cardiac valve formation [24, 30, 41]. Furthermore, we previously demonstrated that *Gata4* is required in the posterior SHF for atrial septation. Both Hedgehog (Hh) signaling and *Pten*-mediated cell-cycle progression were shown to be downstream of *Gata4* in atrial septation [42]. However, the mechanistic requirement for Gata4 in OFT development is less clear. For example, from the multiple *Gata4* transcriptional targets that have been identified in the context of heart development, including *Nppa, α-MHC, α-CA, B-type natriuretic peptide* (*BNP*)*, Ccnd2*, and *Cyclin D2*, and *Mef2c* [20, 23, 24, 26, 43, 44], only *Mef2c* has a functional role in OFT development [12].

In this study, we investigated the mechanistic requirement for *Gata4* in OFT development. We found that *Gata4* heterozygosity in SHF hedgehog (Hh)-receiving cells recapitulates the OFT misalignment observed in *Gata4* germline heterozygotes in mice. Gata4 heterozygous embryos had decreased numbers of SHF-derived cells populating the anterior SHF and the developing OFT at E10.5. By genetic inducible fate mapping (GIFM), Hh-receiving cells fail to migrate properly into the OFT of *Gata4* mutant mice. We have previously reported that Gata4 acts upstream of Hh-signaling for atrial septation [42]. Here we observed more severe OFT defects observed in embryos with SHF-specific heterozygosity of *Gata4* and *Smo*, the obligate Hh signaling receptor. Furthermore, rescue of *Gata4*-mediated OFT misalignment by constitutive activation of Hh-signaling indicated a consistent epistatic relationship between Gata4 and Hh signaling in OFT development. Furthermore, upregulation of *Gata6* in the *Gata4* mutant SHF may provide an explanation for the severity of OFT defects observed in *Gata4* mutant embryos. Our study thereby revealed *Gata4*-dependent pathways contributing to OFT development in *Gata4* heterozygous mouse embryos.

## Results

### GATA4 is required for OFT alignment

*Gata4* is strongly expressed in the heart, pSHF and OFT at E9.5 [27, 42, 50]. There is a gap in expression between the OFT and the pSHF at embryonic day 9.5 (Fig. 1A, indicated by a “↓”).IHC staining for Gata4 at later stages during OFT development showed strong Gata4 expression in the heart, the developing OFT and the pSHF, but only in a limited subset of aSHF cells at E10.5 (Fig. 1B, indicated by a “↓”). At E11.5, both the chamber myocardium and the developing OFT had strong Gata4 expression, however, Gata4 expression was absent from the cardiac neural crest (CNC)-derived distal OFT (Fig. 1C, indicated by a “↓”).

**Figure 1.**
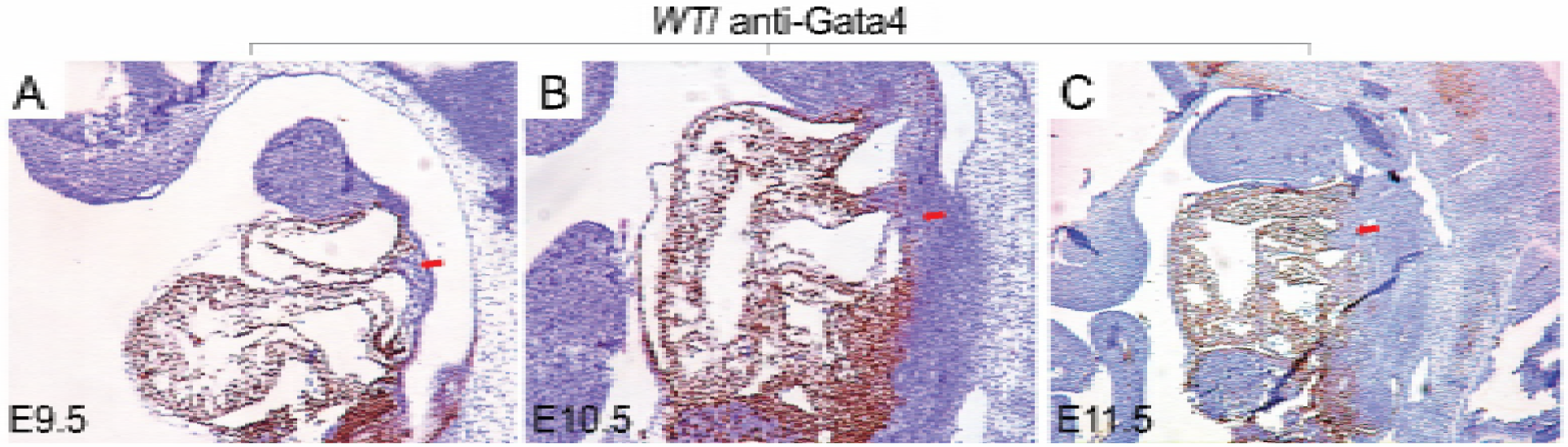
Gata4 is strongly expressed in the developing heart, the OFT and the pSHF. Gata4 expression was detected in *wildtype* mouse embryos by IHC at (A) E9.5, (B) E10.5 and (C) E11.5. Red arrows indicate anterior second heart field at E9.5 or E10.5 (A and B), and proximal outflow tract at E11.5 (C). Magnificence: A: 40X; B and C: 100X

Gata4 was previously reported to be required for OFT alignment [27]. To study the role of Gata4 in OFT development, we re-examined *Gata4* heterozygotes for OFT defects. As described previously [42], *Gata4* heterozygotes were generated by crossing *Gata4^fl/+^* with *Ella^Cre^*, which drives Cre expression in the germline [51] to effect germline *Gata4* deletion. The Gata4 germline deletion was ensured by genotyping using the embryo tail DNA. Whereas *Gata4^fl/+^* (n = 13) and *Ella^Cre/+^* (n = 12) embryos demonstrated normal heart at E14.5 (Figs. 2A and A’, 2B and B’), 61.1% of *Gata4^+/-^*; *Ella^Cre/+^* embryos demonstrated VSD and DORV (Figs. 2C’, 11/18, P=0.0004). Consistent with our prior work, we observed primum ASDs with absence of the DMP in 8/18 *Gata4^+/-^*; *Ella^Cre/+^* embryos [42] (Figs. 2C).

**Figure 2.**
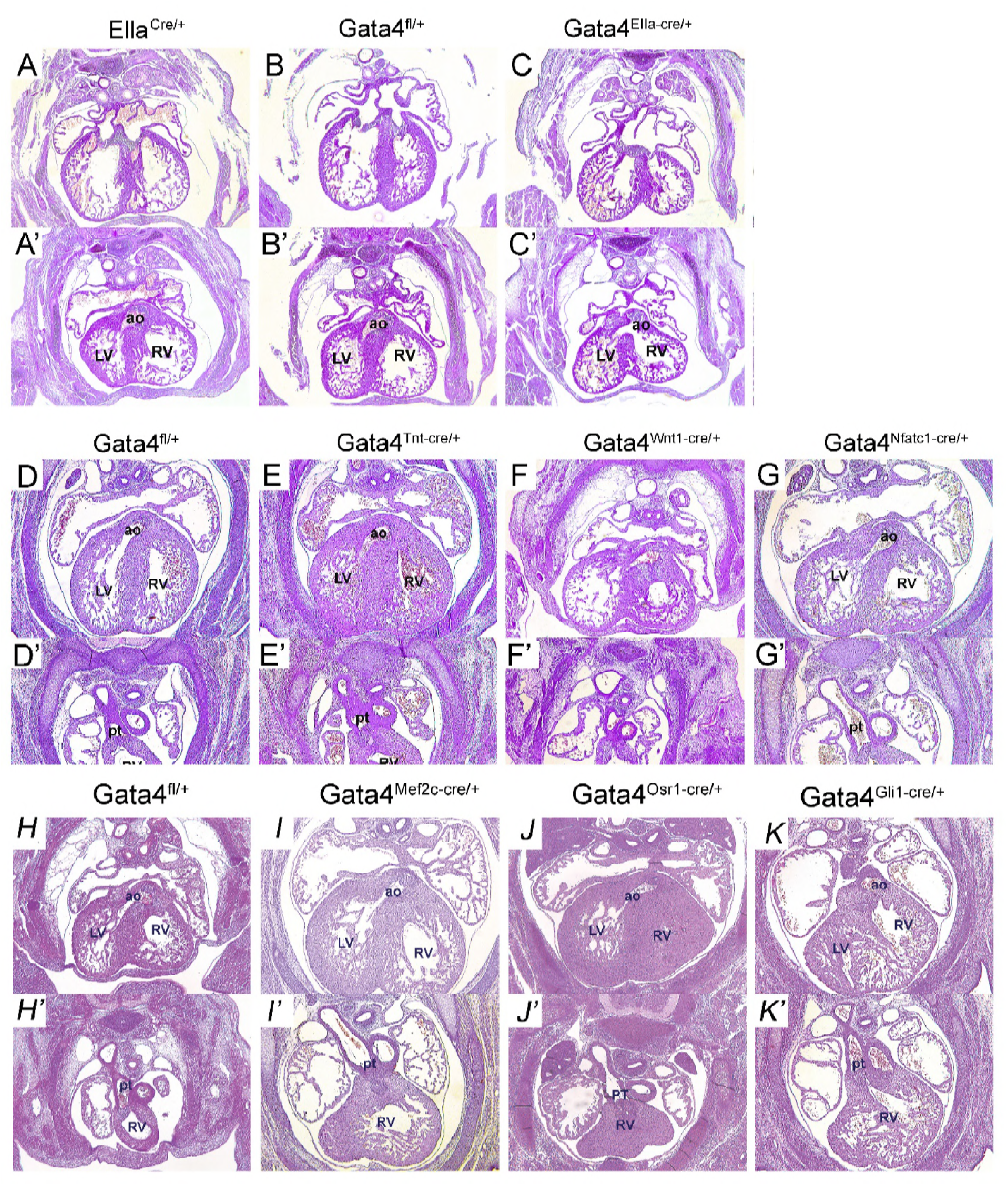
Gata4 is required in Hh-receiving cells for OFT development. (A-G’) Histology of Gata4 transgenic mouse embryo heart at E14.5. Statistics were summarized in table 1. Histology of Gata4 transgenic mouse embryo heart at E13.5. LV, left ventrium; RV, right ventrium; ao, aorta artery, PT, pulmonary trunk. Magnificence: 40X (H-K’) Histology of Gata4 transgenic mouse embryo heart at E14.5. Histology of Gata4 transgenic mouse embryo heart at E13.5. LV, left ventrium; RV, right ventrium; ao, aorta artery, PT, pulmonary trunk.

To determine the lineage requirement for *Gata4* in AV septation, we analyzed mouse embryos haploinsufficient for *Gata4* in the myocardium, CNC, endocardium or SHF. We combined *Tnt: Cre* [52] with *Gata4^fl/+^* to create *Gata4* haploinsufficiency in the myocardium. Normal OFT alignment was observed in all *Tnt^Cre/+^; Gata4^fl/+^* (12/12) and littermate control *Gata4^fl/+^* embryos (9/9) at E13.5 (P=1) (Figs. 2E and E’ vs. 2D and D’, P=1). We combined *Wnt1: Cre* [53, 54] with *Gata4^fl/+^* create *Gata4* haploinsufficiency in the CNC. Normal OFT alignment was observed in all *Wnt1^Cre/+^; Gata4^fl/+^* mutant embryos (24/24) and littermate control *Gata4^fl/+^* embryos (16/16) at E13.5 (Figs. 2F and F’ vs. 2D and D’, P=1). We combined *Nfat1c: Cre* [53, 54] with *Gata4^fl/+^* create *Gata4* haploinsufficiency in the endocardium. Normal OFT alignment was observed in nearly all *Nfatc1^Cre/+^; Gata4^fl/+^* mutant embryos (14/15) and littermate control *Gata4^fl/+^* embryos (10/10) at E13.5 (Figs. 2G and G’ vs. 2D and D’, P=1). These results demonstrated that *Gata4* haploinsufficiency in the myocardium, CNC or endocardium supported normal OFT alignment.

### *Gata4* is required in the SHF Hedgehog (Hh) signal-receiving progenitors for OFT alignment

We hypothesized that Gata4 is required in the aSHF for OFT alignment in aSHF-specific *Gata4* heterozygous mice. We tested this hypothesis by combining *Mef2cAHF: Cre* with *Gata4^fl/+^*. Surprisingly, OFT misalignment with DORV was only observed in 1 out of 22 embryos and none in the littermate controls (Fig. 2I and I’ vs. 2H and H’, P=1). We next tested if Gata4 is required in the pSHF for OFT alignment in in pSHF-specific *Gata4* heterozygous mice by crossing *Osr1 ^CreERT2/+^* [46, 47] with *Gata4^fl/+^*. Similarly, neither *Gata4^fl/+^*; *Osr1 ^CreERT2/+^* embryos (0/5) nor littermate control *Gata4^fl/+^* embryos (0/6) demonstrated OFT misalignments at E14.5 (Fig. 2J and J’ vs. 2H and H’, P=1). These results demonstrated that *Gata4* haploinsufficiency in either aSHF or pSHF supported normal OFT alignment.

Previous studies have shown that SHF Hh signal-receiving progenitors localized in both the aSHF and the pSHF, and regulated the migration of SHF toward the OFT and inflow tract (IFT) to form the pulmonary artery and the atrial septum separately [45, 55, 56]. We combined *Gli1^Cre-ERT2^* with *Gata4^fl/+^* to create *Gata4* haploinsufficiency in SHF Hh signal-receiving progenitors. CreERT2 was activated by tamoxifen (TM) administration at E7.5 and E8.5 in *Gli1^Cre-ERT2^*; *Gata4^fl/+^* embryos. With TM administration at E7.5 and E8.5, 66.7% of *Gli1^Cre-ERT2^*; *Gata4^fl/+^* embryos displayed DORV, while the littermate control *Gata4^fl/+^* embryos displayed normal OFT alignment (Figure 2K and K’ vs. 2H, 2H’, 8/12 vs. 0/15, P=0.0002). We concluded that *Gata4* is required in the SHF Hedgehog (Hh) signal-receiving progenitors for OFT alignment.

### Gata6 was overexpressed in the SHF of the Gata4 heterozygotes

*Gata4* and *Gata6* double mutant embryos display PTA [40]. We examined *Gata6* expression in *Gata4* mutants. Gata6 was expressed in the heart, the OFT and strongly in the splanchnic mesoderm (Fig. 3A, arrow), but not neural crest cell derivatives (Fig. 3A, arrowhead) of the *Gata4^fl/+^* embryo at E9.5. In *Gata4* knockdown embryos specifically in the Hh-receiving cells, *Gata6* expression domain was strongly enhanced in the OFT and the splanchnic mesoderm. Consistently enhanced expression of *Gata6* in the OFT and the SHF of the *Gata4^fl/fl^; Gli1^Cre-ERT2/+^* was further confirmed by the real-time PCR at the mRNA level (Fig. 3B). The *Gata4* expression in the SHF of *Gata4^fl/fl^; Gli1^Cre-ERT2/+^* mouse embryo was 2.7-fold that observed in control *Gata4^fl/+^* embryos (P<0.05). *Gata6* expression in the OFT of the *Gata4^fl/fl^; Gli1^Cre-ERT2/+^* mouse embryo was 4.4-fold that of the littermate control (P<0.01). Our results suggested a negative association between the expression of *Gata4* and *Gata6* in the SHF and developing OFT.

**Figure 3.**
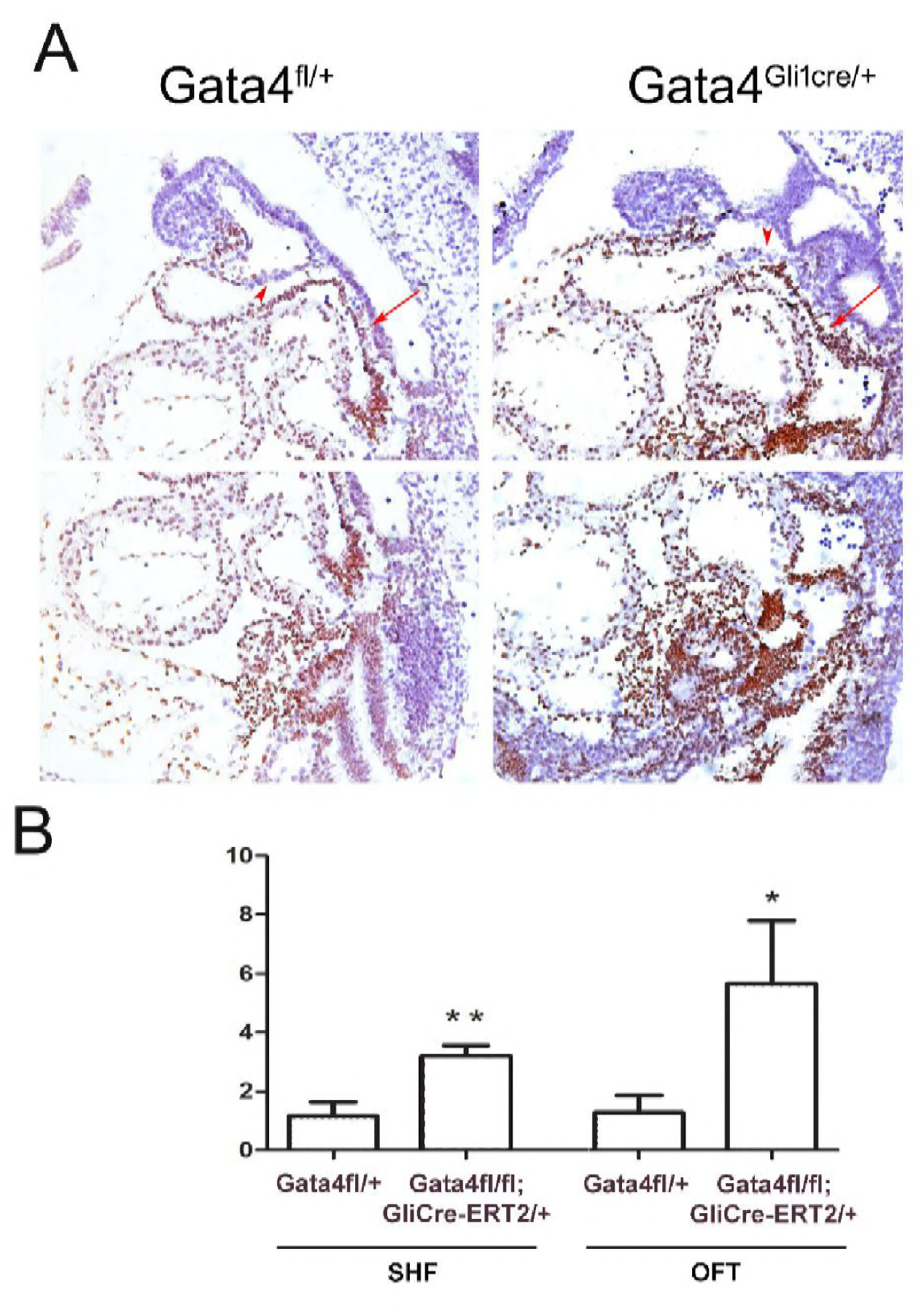
Gata6 was overexpressed in the OFT and the SHF of the Gata4 mutant embryos at E9.5. (A) IHC of the Gata6 in *Gata4^fl/+^* and *Gata4^fl/+^; Gli1^Cre-ERT2/+^* embryos at E9.5. the arrowhead indicated the NCCs-derived cells and the arrow indicates the splanchnic mesoderm. Magnificence: 200X. (B) Gata6 was measured by realtime-PCR in the micro-dissected SHF and the OFT of the *Gata4^fl/+^* and *Gata4^fl/fl^; Gli1^Cre-ERT2/+^* embryos at E9.5. *p<0.1, **p<0.05, n=3

### *Gata4* regulates cell proliferation in the OFT conal cushion

We wonder if Gata4 is required for proliferation during the OFT cushion development. Cell proliferation was examined by BrdU incorporation at E11.5. *Gata4^fl/+^; Gli1^Cre-ERT2/+^* embryos demonstrated 17% fewer BrdU-positive SHF cells in the OFT conal cushion (Fig. 4C vs. 4A and 4E; *P* =0.0134), but not the OFT truncal cushion (Fig. 4D vs. 4B and 4F; *P* =0.1998), compared to the littermate *Gata4^fl/+^*embryos at E11.5. This result demonstrate that *Gata4* is required for normal cell proliferation in OFT conal cushion development. We assessed cell death by TUNEL staining and observed no differences in either the conal or truncal cushion between *Gata4fl^/+^; Gli1^Cre-ERT2/+^*and the *Gata4^fl/+^*embryos (Fig. 4G - 4J). Together, these findings define a requirement for *Gata4* in the proliferation but not in the survival of OFT conal cushion cells.

**Figure 4.**
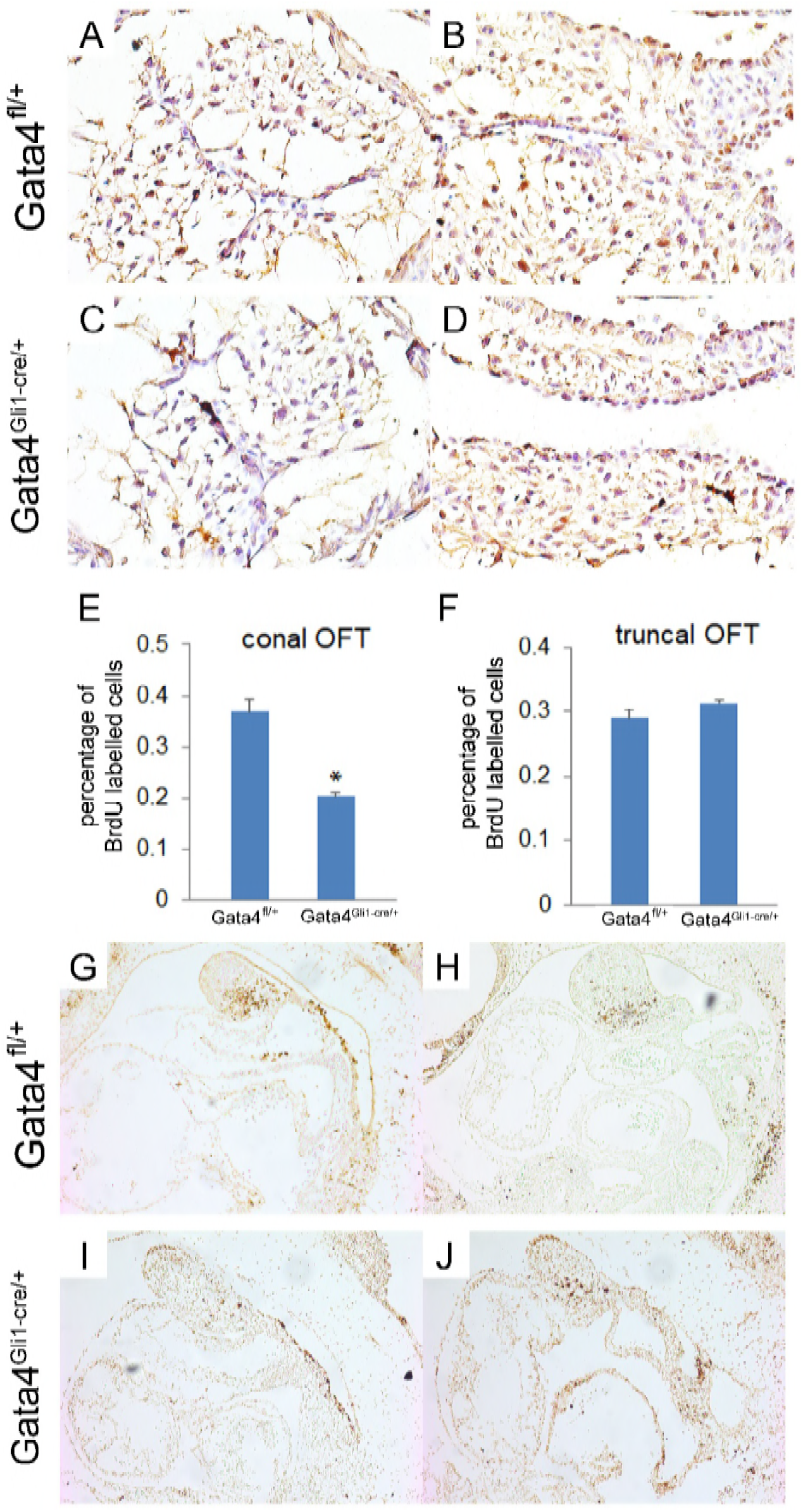
Gata4 regulates cell proliferation in conal OFT. (A-D) BrdU staining in conal OFT and truncal OFT in *Gata4^fl/+^; Gli1^Cre-ERT2/+^* embryos and control embryos at E10.5. Magnificence: 400X. (E and F) Quantification of BrdU labelled cells. Data is presented as mean±SE, *p<0.05, n=3, One-way ANOVA. (G-J) TUNEL staining in both *Gata4^fl/+^; Gli1^Cre-ERT2/+^* embryos and control embryos at E10.5. Magnificence: 100X

### Rescue of SHF proliferation by disruption of *Pten* does not rescue DORV in *Gata4* mutant embryos

Our previous study demonstrated that *Gata4* regulates the cell cycle progression in posterior SHF cardiac precursors and that genetically targeted disruption of *Pten* rescued the proliferation defects in SHF of the *Gata4* heterozygotes [57]. Hence, we examined whether proliferation rescue in SHF, by *Pten* downregulation (TMX at E7.5 and E8.5), could rescue DORV in Hh-receiving cell-specific *Gata4* heterozygotes. We observed that decreased *Pten* dose caused only one DORV, but no ASD, in 20 embryos (Fig. 5A-C). Consistent with our previous report, although the ASD in *Gli1^Cre-ERT2/+^;Gata4^fl/+^* embryos was rescued by *Pten* downregulation (Fig. 5C vs. 5B, 1/20 in *Gli1^Cre-ERT2/+^;Gata4^fl/+^;Pten^fl/+^* vs. *14/29 in Gli1^Cre-ERT2/+^*;Gata4*^fl/+^*, P = 0.0013), *Gli1^Cre-ERT2/+^*;Gata4*^fl/+^*;Pten*^fl/+^* embryos still displayed DORV with an incidence rate unchanged from *Gli1^Cre-ERT2/+^*;*Gata4^fl/+^* embryos (Fig. 5E vs. 5F, 12/29 vs. 6/20, Table 1, P = 0.5495). This data suggested to us that correction of the SHF proliferation defects was not able to rescue the OFT misalignment of the *Gata4* mutant embryos.

**Figure 5.**
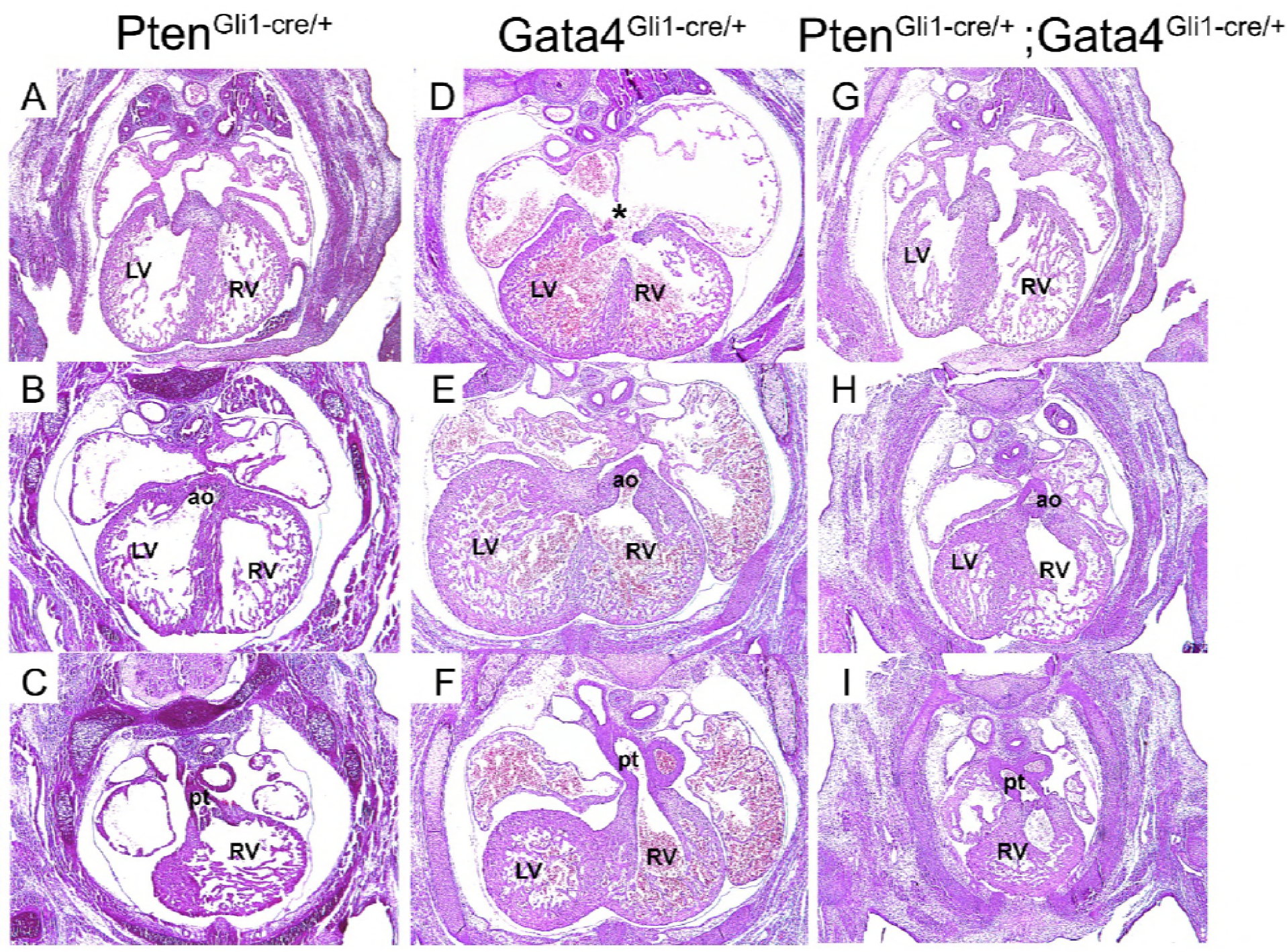
Genetically targeted ablation of Pten rescues atrioventricular septal defect. (A-I) Histology of Gata4 transgenic mouse embryo heart at E13.5. LV, left ventrium; RV, right ventrium; ao, aorta artery, PT, pulmonary trunk. Magnificence: 40X.

**Table 1.**
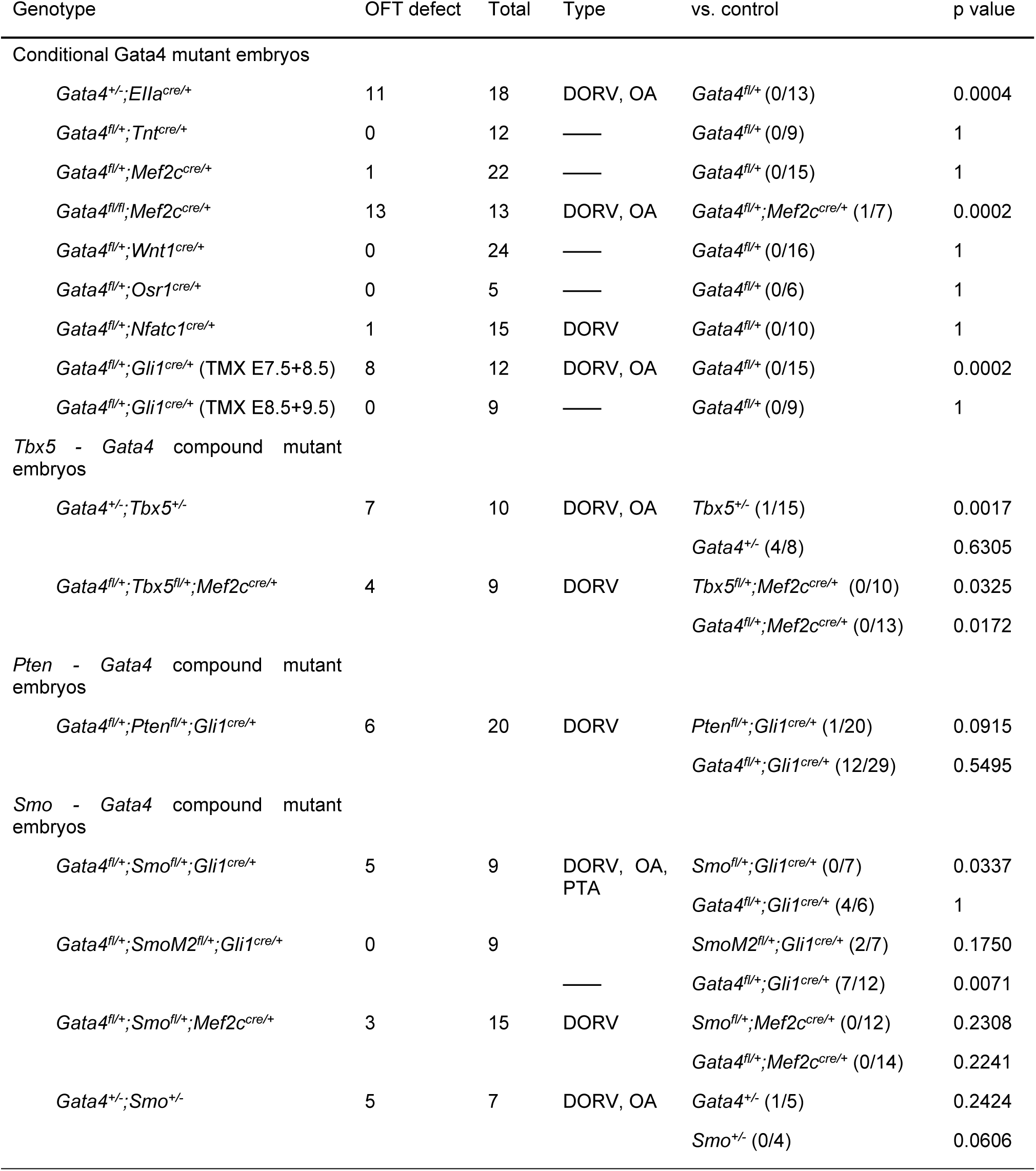
Incidence of OFT defect in Gata4 mutant embryos

### Gata4 acts upstream of Hh signaling in OFT development

We have previously reported that *Gata4* acts upstream of Hh-signaling for atrial septation [42]. The requirement of *Gata4* in *Hh-*receiving cells for OFT alignment suggested that *Gata4* and *Hh* signaling may interact genetically in the SHF for OFT development. We tested this hypothesis in the *Gata4* and *Smo* compound heterozygotes (*Gata4^fl/+^;Smo^fl/+^;Gli1^Cre-ERT2/+^*) versus littermate controls (*Gata4^fl/+^; Gli1^Cre-ERT2/+^* or *Smo^fl/+^;Gli1^Cre-ERT2/+^*). Consistent OFT defects were observed in compound *Gata4; Smo* embryos (*Gata4^fl/+^;Smo^fl/+^;Gli1^Cre-ERT2/+^*) (5/9, Fig 6C - 6E) whereas no OFT defects were observed in *Smo^fl/+^;Gli1^Cre-ERT2/+^*embryos (0/7, Fig 6B and B’; P= 0.0337). The total incidence of OFT defects occured in the *Gata4^fl/+^;Smo^fl/+^;Gli1^Cre-ERT2/+^* was not different than in the *Gata4^fl/+^; Gli1^Cre-ERT2/+^* embryos (Fig 6C-E, 5/9 vs. 4/6, P= 0.7326). However, more severe range of OFT defects was observed in *Gata4^fl/+^;Smo^fl/+^;Gli1^Cre-ERT2/+^* embryos, including DORV (3 out of 5, Figs. 6C and C’), OA (1 out of 5, Figs. 6D and D’) and persistent truncus arteriosus (PTA) (1 out of 5, Figs. 6E and E’). PTA, caused by a combined defect of alignment and separation, was only observed in *Gata4^fl/+^;Smo^fl/+^;Gli1^Cre-ERT2/+^*. This result suggest an interaction between *Gata4* and Hh-signaling in OFT development.

**Figure 6.**
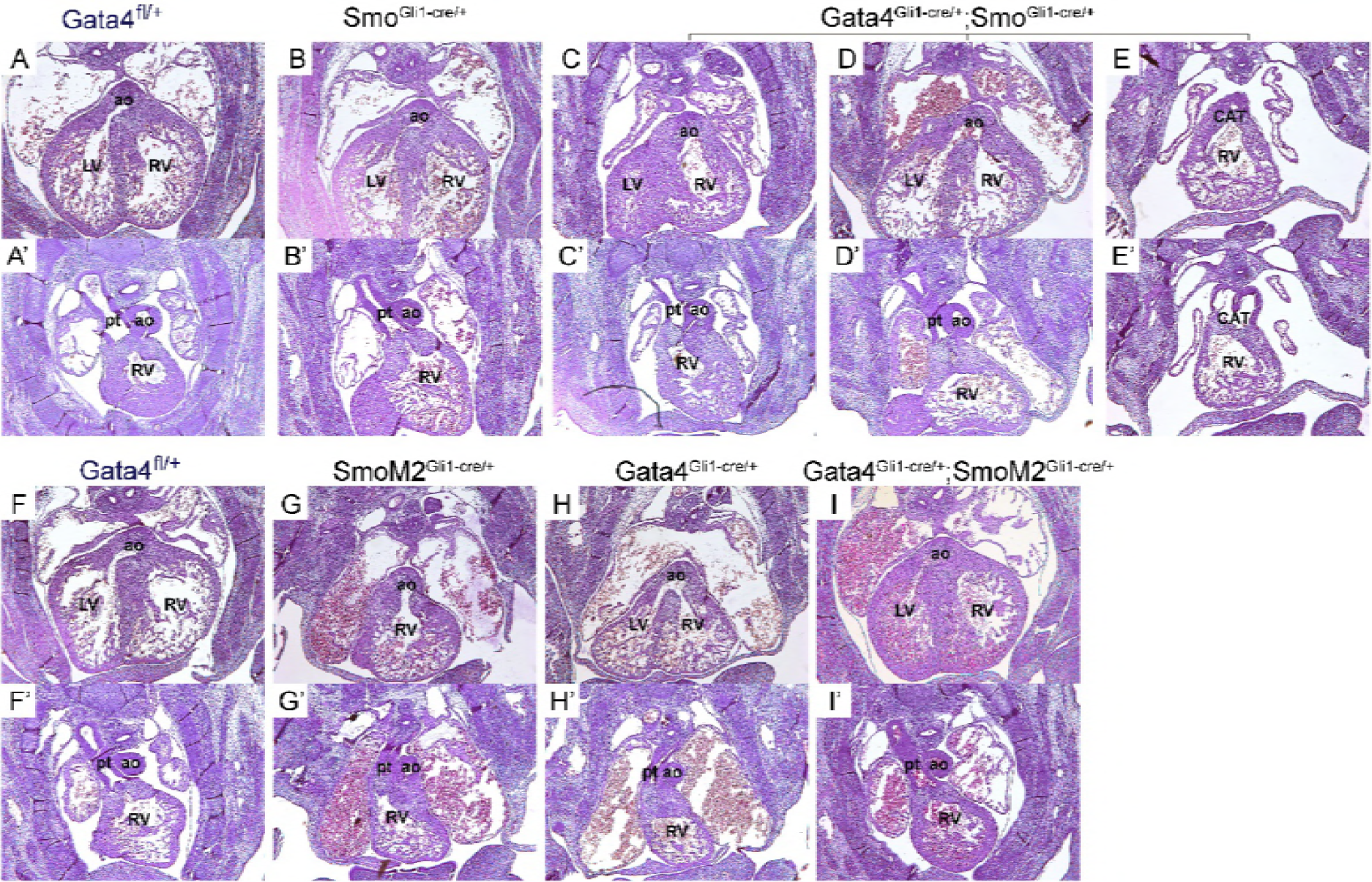
Gata4 acts upstream of Hh signaling pathway. (A-I’) Histology of Gata4 transgenic mouse embryo heart at E14.5. LV, left ventrium; RV, right ventrium; ao, aorta artery, PT, pulmonary trunk; CAT, common artery trunk. Magnificence: 40X.

We tested the hypothesis that *Gata4* actis upstream of Hh-signaling for OFT development using a genetic epistasis study. We tested whether increased Hh-signaling via a constitutively activated Smo mutant, *SmoM2* [58], could rescue the OFT misalignment in *Gata4*-heterozygotes. DORV was observed in 28.6% of littermate control *Gli1^Cre-ERT2/+^;R26-SmoM2^fl/+^*embryos (2/7) (Fig. 6G and G’) and 58.3% of littermate control *Gli1^Cre-ERT2/+^;Gata4^fl/+^*embryos at E14.5 (7/12) (Fig. 6H and H’). In contrast, none of *Gata4^fl/+^;Gli1^Cre-ERT2/+^;R26-SmoM2^fl/+^* embryos showed DORV (Fig. 6I and I’), indicating significant rescue by *R26-SmoM2^fl/+^*, *Gli1^Cre-ERT2/+^*(Fig. 6I vs Fig. 6H, P = 0.0071, Table 1). This results demonstrated rescue of DORV in *Gata4*-mutant embryos by constitutive Hh signaling.

### *Gata4* is required for the contribution of Hh-receiving cells to the OFT

Hh signaling has been reported to regulate the migration of SHF Hh-receiving cells toward the arterial pole of the heart [45]. We therefore hypothesized that *Gata4* is required for the SHF Hh-receiving cells migration toward the developing OFT. We tested this hypothesis using genetic inducible fate mapping (GIFM) [59]. The Hh-receiving lineage cells were marked in *R26R^fl/+^;Gli1^Cre-ERT2/+^*embryos by TM administration at E7.5 and E8.5 and *β-gal* expression was evaluated at E10.5 in Gata4 heterozygotes. The total number of *β-gal* positive cells was obtained by counting those on each individual sections and adding up all through the SHF and the OFT. We have previously reported decreased number of Hh-receiving cells in the pSHF at E9.5 associated with developing defects of DMP in the *Gata4^fl/+^;R26R^fl/+^;Gli1^Cre-ERT2/+^*embryos [57]. We observed that there were also significantly less Hh-receiving cells within the aSHF region (Fig. 7A vs. 7D and Fig. 7G, 334.0 ± 1.4 vs. 186.7 ± 4.9, P=0.009) of the *Gata4^fl/+^;R26R^fl/+^;Gli1^Cre-ERT2/+^*embryos. The cells of Hh-receiving lineage were observed in the developing OFT at this stage. By counting the number of β-galactosidase-expressing cells in the proximal half (Fig. 7B vs. 7E and 7H, 49.7 ± 9.6 vs. 26.7 ± 6.7, P=0.097) and the distal half of the OFT (Fig.7C vs. 7F and 7I, 91.7 ± 9.2 vs. 57.0 ± 1.4, P=0.0362), we found that both of the regions of the *Gata4* heterozygotes had less β-galactosidase-expressing cells than the littermate controls (Figs. 7E and 7F).

**Figure 7.**
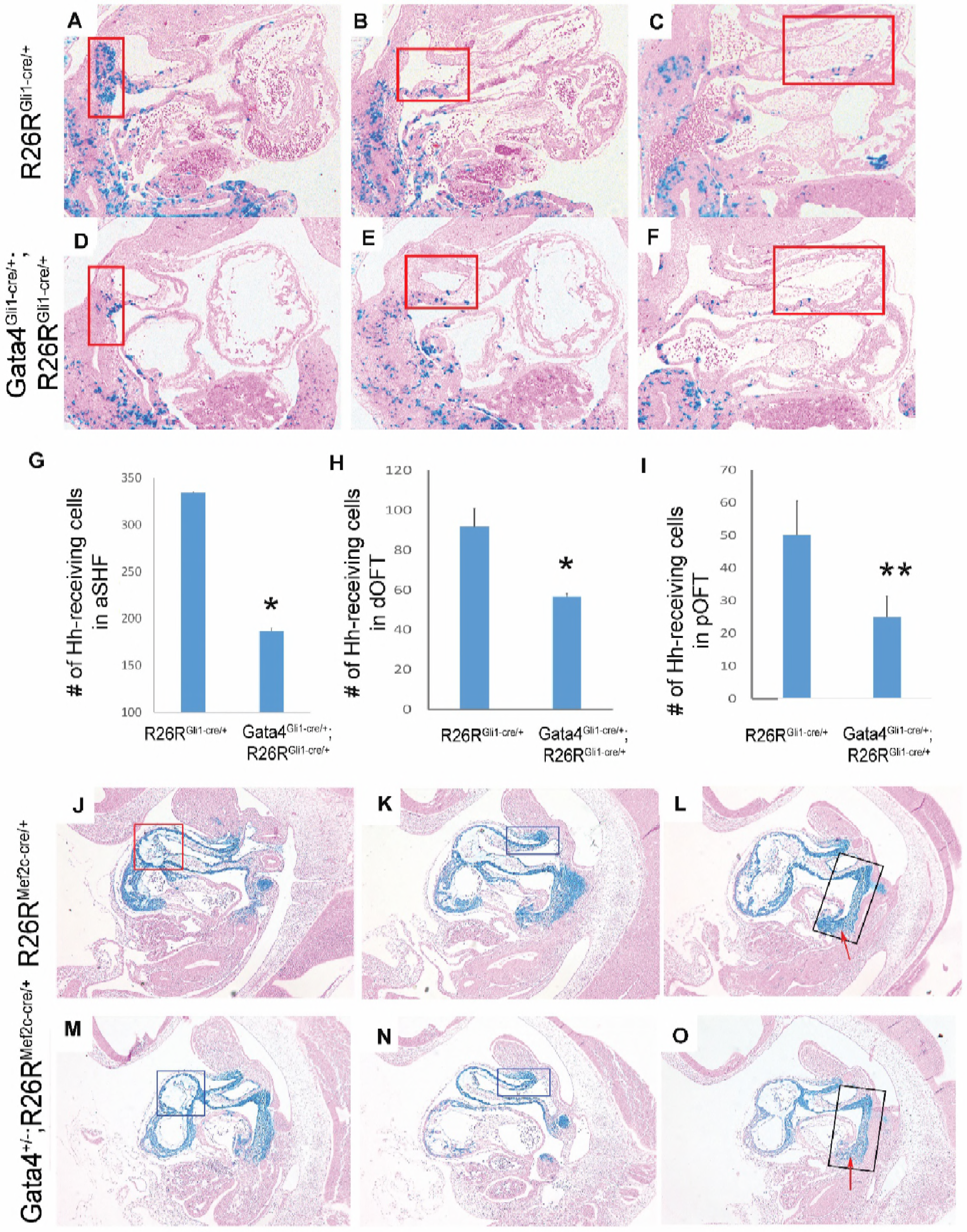
*Gata4* is required for the contribution of Hh-receiving cells to the OFT. (A-F) LacZ staining of Gli1-expressing cells in Gata4 transgenic mouse embryos at E10.5 focusing on aSHF (E and H), dOFT (F, I) and pOFT (G, J). (G-I) Quantification of stained cells within selected regions. Data is presented as mean±SE, *p<0.05, ** p<0.1, n=3, One-way ANOVA. (J-O) LacZ staining of cells with Mef2cAHF:Cre expression in Gata4 transgenic mouse embryos at E10.5. The red arrow indicated a developing DMP region. Magnificence: A-D and A’-D’ 40X; E-J: 100X; N-S: 100X

To examine if Gata4 haploinsufficency influenced the SHF cell recruitment within the proximal OFT, we analyzed the fate map of SHF lineage cells in the OFT of the *Gata4* heterozygotes. Defined by *Mef2cAHF:Cre* expression: β-galactosidase-expressing cells, the total number of the SHF lineage cells within the proximal half and the distal half of the OFT were compared between the *Mef2cAHF::Cre;Gata4^fl/+^; R24R^fl/+^* and the *Mef2cAHF::Cre;R24R^fl/+^*embryos at E10. The number of SHF lineage cells populating the proximal OFT of the *Mef2cAHF::Cre;Gata4^+/-^; R24R^fl/+^* embryos was significantly less than that those in control *Mef2cAHF::Cre; R24R^fl/+^* embryos (Fig. 7J vs. 7M); however, this decrement was not observed in the distal OFT (Fig. 7K vs. 7N). The distribution pattern of the SHF lineage was not different in the *Mef2cAHF::Cre;Gata4^+/-^; R24R^fl/+^* and the *Mef2cAHF::Cre;R24R^fl/+^*embryos (Figs. 7L vs. 7O). AS a control, we observed fewer cells populating the developing dorsal mesocardium protrusion (DMP) in *Mef2cAHF::Cre;Gata4^+/-^; R24R^fl/+^*(red arrow, Fig. 7L vs. 7O), consistent with our previous report that *Gata4* is required in the SHF for the DMP [42]. These results demonstrated the requirement of *Gata4* for the SHF lineage cells populating in the developing OFT.

## Discussion

The requirement of Gata4 for OFT development has been reported in mice and human, and mouse Gata4 mutations cause DORV [22, 27, 40]. Here we demonstrate that Gata4 is required in the SHF Hh-receiving cells for OFT alignment in the SHF. Our previous study has demonstrated that Gata4 is required for Hh signaling in the SHF for cell proliferation. However, the current study suggested that the cell proliferation defects in the SHF caused by Gata4 mutation may not directly associate with the OFT misalignment; instead, the migration defects of the SHF cells is. And the migration defects were associated with disrupted Hh-signaling, because the OFT misalignment was rescued by over-activating of Hh-signaling. In addition, our data suggested breaking down the threshold of GATA including *Gata4* and *Gata6*, and Hh signaling tone might be associated with the severity of OFT defects.

The SHF was initially described as a progenitor field for the cardiac OFT and a rich literature has established the requirement of anterior SHF contributions for OFT development [5, 10-19, 60-63]. More recently, the contribution of posterior SHF cardiac progenitors to the OFT and the future subpulmonary myocardium has been reported, however, the mechanistic requirement for this contribution is not well understood [45, 64-66]. The cell lineage in which Gata4 is required for OFT development has not been reported. Gata4 is expressed in both the aSHF and pSHF, although its expression is much stronger in the pSHF than in the aSHF [57]. The decreased number of *Mef2C-AHF::Cre* positive cells in the proximal OFT cushion of E10.5 *Gata4*^−^*^/+^* embryos demonstrated that Gata4 plays a role in adding the SHF progenitor cells to the developing OFT. However, surprisingly, OFT defects were not observed in either aSHF-specific or pSHF-specific Gata4 happloinsufficiency. Instead, we found that OFT defects severity and incidence rate in embryos with *Gata4* haploinsufficienc in *Hh*-receiving cells were identical to those in *Gata4*^−^*^/+^* embryos. Because Hh-receiving cells are located throughout the SHF, these observations suggest Gata4 is required in both pSHF and aSHF progenitor cells for OFT alignment.

We provided evidence that Gata4 acts upstream of Hh-signaling in the SHF for OFT development. The *Gata4*^−^*^/+^* embryos have combined phenotypes of ASD and DORV [57]. We previously reported the Gata4-Hh-signaling regulation in atrial septation and identified *Gli1* as the direct target of GATA4 [42]. Here, our data of less percentile of BrdU+ cells in the conal cushion of the OFT at E11.5 of the *Gata4^fl/+^*; *Gli1^Cre-ERT2/+^* embryos, suggesting a role of Gata4 in regulating the OFT cushion cell proliferation. In the posterior SHF, Gata4-Hh-signaling controls cell cycle progression and thereby the proliferation of the cardiac progenitors. Diminished Gata4-Hh signaling causes a failure of development of the DMP, the anlage of the atrial septum, resulting in ASDs [57]. The effect of this pathway on the cell cycle is balanced by Pten via transcriptional inhibition of Cyclin D4 and Cdk4 [20, 57], as DMP hypoplasia and SHF cell cycle defects are rescued by *Pten* knockdown [57]. In the current study, *Pten* knockdown was unable to rescue DORV or OA defects in *Gata4* heterozygous mutants. This observation suggests that correction of SHF cell proliferation is not sufficient to support a normal OFT development in Gata4 mutants, and that Gata4 plays a distinct role in the anterior SHF.

Endodermal Hh signaling is required for the survival of the pharyngeal endoderm, which cell non-autonomously affects SHF survival and OFT lengthening [55]. In our study, increased apoptosis was not observed in the SHF of *Gata4* heterozygote mutant embryos [57]. However, fate mapping of the SHF using either *Mef2c::Cre* or the *Gli1Cre:ERT2* disclosed less SHF-derived cells in the distal OFT in Gata4 mutant embryos. Specifically, there was decreased number of SHF Hh-receiving cells throughout the migration route from the SHF into the OFT: from the dorsal mesocardium through the rostral splanchnic mesoderm, past the distal OFT to the proximal OFT. Hh-receiving progenitors have been found to migrate from the aSHF to populate the pulmonary trunk between E9.5 to E11.5 [45], suggesting that Hh-signaling is required for SHF cell migration. The observation that DORV in Gata4 mutant embryos can be rescued by constitutive Hh-signaling implies correction of both the proliferation and the migration defects of the SHF cardiac progenitors, not proliferation defects only. Overall, here we provide cellular, molecular and genetic evidence that Gata4-Hh signaling hierarchy is required in OFT alignment, with specific regulation of both proliferation and migration of SHF progenitors.

Although important *Gata4* transcriptional targets in the heart have been identified [20, 26, 44], *Gata4*-dependent molecular pathways required for OFT development have remained unknown. We previously identified Gli1 as a downstream target of Gata4 in the posterior SHF for atrial septation [42]. In the current study we further demonstrated that Gata4 regulated Hh-signaling via transcriptional regulation through *Gli1* in the anterior SHF for cell migration and OFT alignment. In addition, we provide evidence that *Gata6* expression is negatively regulated by *Gata4* in the OFT. Enhanced *Gata6* expression in *Gata4* mutants might illustrate a compensatory feedback loop, given that *Gata6* and *Gata4* are redundant for cardiac myocyte differentiation [67, 68]. *Gata4/Gata6* compound heterozygotes displayed persistnat truncus ateriosus (PTA), a severe OFT defect caused by combined alignment and OFT septation defects (40). Here we find that *Gata4/Smo* compound heterozygotes show a similar phenotype. *Gata4* heterozygotes alone do not display PTA, which might be due to the partial recovery of GATA function from enhanced *Gata6* expression. Together with previous study [40], these data suggest a threshold of *Gata4, Gata6*, and *Hh* signaling and that is required for OFT development. This suggests that GATA TFs may be essential for the quantitative regulation of Hh signaling, and that strongly diminished GATA function or diminished GATA and Hh signaling together may cause worse OFT defects through regulation of OFT Hh signaling. Future studies will focus on the quantitative relationship between GATA tone and Hh signaling tone and on the Gata4 dependent gene regulatory network (GRN) [69] for OFT development.

## Materials and methods

### Mouse lines

All mouse experiments were performed in a mixed B6/129/SvEv background. *Gata4^fl/+^, Gli1^CreERT2/+^*, *Mef2cAHF::Cre*, *Tie2^Cre/+^*, *Smo^fl/+^* mouse lines were kind gifts from Dr. Ivan Moskowitz lab (University of Chicago, Chicago)*. TnT^Cre/+^* mouse line was from Dr. Yiping Chen lab (Tulane University, New Orleans). *Nfat1c^Cre/+^* mouse line was from Dr. Bin Zhou lab (Albert Einstein College of Medicine, Bronx, NY). The *SmoM2^fl/+^*, *Osr1^Cre-ERT2/+^* and *EIIa*^cre/+^mouse lines were purchased from the Jackson Laboratory. Mouse experiments were completed according to a protocol reviewed and approved by the Institutional Animal Care and Use Committee of the Texas A&M University and the University of North Dakota, in compliance with the USA Public Health Service Policy on Humane Care and Use of Laboratory Animals.

### Tamoxifen administration and X-gal staining

Tamoxifen (TM) -induced activation of *CreERT2* was accomplished by oral gavage with two doses of 75 mg/kg TM at E7.5 and E8.5 [45, 46]. X-gal staining of embryos was performed as described [45].

### BrdU incorporation and Immunohistochemistry Staining (IHC)

Standard procedures were used for histology and IHC. IHC was performed using the following antibodies: anti-Gata4 (Abcam), anti-Gata6 (Abcam). For BrdU incorporation, pregnant mice were given 100mg BrdU per kg bodyweight at 10mg/mL concentration solutions at E11.25 with two doses, 3 hours and 6 hours before sacrifice, respectively. The BrdU staining was performed using a BrdU In-Situ detection kit (EMD Millipore). For TUNNEL staining, an ApopTag plus peroxidase In-Situ apoptosis detection kit was used (EMD Millipore).

### Micro-dissection of pSHF and RNA extraction

To obtain the pSHF splanchnic mesoderm for use in quantitative realtime-PCR, E9.5 embryos were dissected as described before [47, 48]. The heart, aSHF, and pSHF were collected separately in RNA-later, and then stored at −20°C until genotyping was completed.

### Realtime-PCR

Total RNA was extracted from the PSHF regions of mouse embryos hearts using RNeasy Mini Kit (QIAGEN), according to the manufacturer’s instructions. Two hundred ng of total RNA was reverse transcribed using a SuperScript™ III Reverse Transcriptase kit from Invitrogen. qPCR was performed using a POWER SYBER Green PCR mater mix from Applied Biosystems. Results were analyzed using the delta-delta Ct method with *GAPDH* as a normalization control [49].

## Acknowledgements

We would specifically like to acknowledge the support of Dr. Boon Chew for the study.

